# CosR is a repressor of compatible solute biosynthesis and transporter systems

**DOI:** 10.1101/845297

**Authors:** Gwendolyn J. Gregory, Daniel P. Morreale, E. Fidelma Boyd

**Affiliations:** Department of Biological Sciences, University of Delaware, Newark, DE, 19716

## Abstract

Bacteria accumulate small, organic compounds, called compatible solutes, via uptake from the environment or biosynthesis from available precursors to maintain the turgor pressure of the cell in response to osmotic stress. *Vibrio parahaemolyticus* has biosynthesis pathways for the compatible solutes ectoine (*ectABCasp_ect)* and glycine betaine (*betIBAproXWV)*, four betaine-carnitine-choline transporters (*bcct1-bcct4*) and a second ProU transporter (*proVWX).* Most of these systems are induced in high salt. CosR, a MarR-type regulator, which is divergently transcribed from *bcct3*, was previously shown to be a direct repressor of *ectABCasp_ect* in *Vibrio* species. In this study, we investigated the role of CosR in glycine betaine biosynthesis and compatible solute transporter gene regulation. Expression analyses demonstrated that *betIBAproXWV*, *bcct1*, *bcct3*, and *proVWX* are repressed in low salinity. Examination of an in-frame *cosR* deletion mutant shows induced expression of these systems in the mutant at low salinity compared to wild-type. DNA binding assays demonstrate that purified CosR binds directly to the regulatory region of each system. In *Escherichia coli* GFP reporter assays, we demonstrate that CosR directly represses transcription of *betIBAproXWV*, *bcct3*, and *proVWX*. Similar to *V. harveyi*, we show *betIBAproXWV* is positively regulated by the LuxR homolog OpaR. Bioinformatics analysis demonstrates that CosR is widespread within the genus, present in over 50 species. In several species, the *cosR* homolog was clustered with the *betIBAproXWV* operon, which again suggests the importance of this regulator in glycine betaine biosynthesis. Incidentally, in four *Aliivibrio* species that contain ectoine biosynthesis genes, we identified another MarR-type regulator, *ectR*, clustered with these genes, which suggests the presence of a novel ectoine regulator. Homologs of EctR in this genomic context were present in *A. fischeri, A. finisterrensis, A. sifiae* and *A. wodanis*.

**Importance:** *Vibrio parahaemolyticus* can accumulate compatible solutes via biosynthesis and transport, which allow the cell to survive in high salinity conditions. There is little need for compatible solutes under low salinity conditions, and biosynthesis and transporter systems are repressed. However, the mechanism of this repression is not fully elucidated. CosR plays a major role in the repression of multiple compatible solute systems in *V. parahaemolyticus* as a direct negative regulator of ectoine and glycine betaine biosynthesis systems and four transporters. Homology analysis suggests that CosR functions in this manner in many other *Vibrio* species. In *Aliivibrio* species, we identified a new MarR family regulator EctR that clusters with the ectoine biosynthesis genes.

## Introduction

Halophilic bacteria such as *Vibrio parahaemolyticus* encounter a range of osmolarities in the environment. To combat the loss of turgor pressure due to efflux of water in high osmolarity conditions, bacteria have developed a strategy that involves the accumulation of compatible solutes in the cell (1–3). Compatible solutes, as the name suggests, are organic compounds that are compatible with the molecular machinery and processes of the cell, and include compounds such as ectoine, glycine betaine, trehalose, glycerol, proline, glutamate, and carnitine, among others (1, 4–9). Compatible solutes are taken up from the environment or synthesized from various precursors in response to osmotic stress, which allows cells to continue to grow and divide even in unfavorable environments (2, 4, 10, 11).

*Vibrio parahaemolyticus* possesses compatible solute biosynthesis pathways for ectoine and glycine betaine (12). Ectoine biosynthesis is *de novo* in *V. parahaemolyticus*, requiring aspartic acid as the precursor, which can be supplied by the cell (13). Aspartic acid is converted to ectoine by four enzymes, EctA, EctB, EctC and Asp_Ect, encoded by the operon *ectABCasp_ect* (14). Ectoine biosynthesis begins with L-aspartate-β-semialdehyde, which is also pivotal to bacterial amino acid and cell wall synthesis (14). Asp_Ect is a specialized aspartokinase dedicated to the ectoine pathway that, among Proteobacteria, is present only in alpha, gamma and delta species (15). Searches of the genome database demonstrated that ectoine biosynthesis genes are present in nearly 500 species. Of these, nearly a third also produce 5-hydroxyectoine by the action of an additional gene product, ectoine hydroxylase, encoded by *ectD* (16). A recent study has shown that in *V. parahaemolyticus* the quorum sensing response regulator OpaR is a negative regulator of *ect* gene expression (17). It was also shown that in this species, similar to *V. cholerae*, a multiple antibiotic resistance (MarR)-type regulator named CosR is a repressor of *ectABCasp_ect* (17, 18).

Production of glycine betaine takes place in a two-step oxidation from the precursor choline, which is acquired exogenously. *De novo* biosynthesis of glycine betaine has been identified in only a few species of halophilic bacteria (19–24). The two-step oxidation proceeds with choline conversion to glycine betaine by the products of two genes *betB* and *betA*, which encode betaine-aldehyde dehydrogenase and choline dehydrogenase, respectively (25, 26). In *E. coli*, these genes are encoded by the operon *betIBA*, with the regulator BetI shown to repress its own operon (27, 28). In all *Vibrio* species that biosynthesize glycine betaine, the *betIBA* genes are in an operon with the *proXWV* genes, which encode a ProU transporter (12, 13, 29). Regulation of glycine betaine biosynthesis has been studied in several species, but few direct mechanisms of regulation have been shown beyond BetI (27, 28, 30–33). In *V. harveyi*, a close relative of *V. parahaemolyticus*, *betIBAproXWV* was shown to be positively regulated by the quorum sensing master regulator LuxR (32–33).

It is energetically favorable to the cell to uptake compatible solutes from the environment rather than to biosynthesize them, and Bacteria and Archaea encode multiple osmoregulated transporters (9, 34–39). ATP-binding cassette (ABC) transporters are utilized to import exogenous compatible solutes into the cell and include ProU (encoded by *proVWX*) in *E. coli* and *Pseudomonas syringae*, OpuA in *Lactococcus lactis* and *B. subtilis*, and OpuC in *P. syringae* (39–44). *V. parahaemolyticus* encodes two putative ProU transporters of the ABC transporter family, one on each chromosome. ProU1 is encoded on chromosome 1 by *proVWX* (VP1726-VP1728) and ProU2 is encoded on chromosome 2 by the *betIBAproXWV* operon (VPA1109-VPA1114) (12). ProU1 is a homolog of the *E. coli* K-12 ProU, which in this species was shown to bind glycine betaine with high affinity (41, 45, 46). ProU2 is a homolog of the *P. syringae proVXW* (12).

The betaine-carnitine-choline transporters (BCCTs) are single component transporters, the first of which, BetT, discovered in *E. coli*, was shown to transport choline with high-affinity and is divergently transcribed from *betIBA* (47, 48). *Vibrio parahaemolyticus* encodes four BCCTs, three, BCCT1-BCCT3 (VP1456, VP1723, VP1905), on chromosome 1 and one, BCCT4 (VPA0356), on chromosome 2 (12). This is a typical complement of *bcct* genes present among members of the Campbellii clade, which includes *V. alginolyticus, V. campbellii, V. harveyi* and *V. parahaemolyticus*, amongst others (Naughton et al., 2009). The *bcct2* (VP1723) gene is the only *bcct* gene that is not induced by high salinity in *V. parahaemolyticus* (13). All four BCCT transporter were shown to transport glycine betaine amongst others (29). A study in *V. cholerae* demonstrated that a *bcct3* homolog is repressed by the regulator CosR and deletion of the *cosR* gene also affected biofilm formation and motility in this species (18).

In this study, we examined the broader role of CosR in the regulation of glycine betaine biosynthesis and compatible solute transport gene expression in *V. parahaemolyticus*. First, we examined expression of genes encoding osmotic stress response systems in low salinity and used quantitative real-time PCR to determine expression of these genes in a Δ*cosR* deletion mutant. We then determined whether CosR was a direct regulator using DNA binding assays and an *E. coli* plasmid-based reporter assay. We also examined whether *betIBAproXWV* was under the control of the LuxR homolog OpaR in *V. parahaemolyticus*, similar to what was shown in *V. harveyi*. We investigated the distribution of CosR and its genome context among *Vibrionaceae*. Our data indicate that CosR is a key regulator of the osmotic stress response in *V. parahaemolyticus* under low salinity conditions. Distribution of CosR is widespread, and similar genomic context suggests CosR repression of compatible solutes is common among *Vibrio*.

## Results

### Compatible solute biosynthesis and transport genes are downregulated in low salinity

We have previously shown that *V. parahaemolyticus* does not produce compatible solutes ectoine and glycine betaine during growth in minimal media (M9G) supplemented with 1% NaCl (M9G1%) (12, 13). Here we quantified expression levels of both biosynthesis operons in M9G1% or M9G3%. RNA was isolated from exponentially growing wild-type *V. parahaemolyticus* RIMD2210633 cells, at optical density 595 nm (OD_595_) 0.45, after growth in M9G1% or M9G3%. Real time quantitative PCR (qPCR) was performed to determine relative expression levels. Expression analysis shows that ectoine biosynthesis genes *ectA* and *asp_ect* are differentially expressed in M9G1% as compared to expression in M9G3%. *ectA* is significantly downregulated 794.6-fold and *asp_ect* is significantly downregulated 204.9-fold in M9G1% (Fig. 1A). The *betIBAproXWV* operon is also significantly repressed in M9G1%, with fold changes of 25.8-fold, 22-fold, 33.7-fold, and 52.8-fold for *betI*, *betB*, *proX*, and *proW*, respectively (Fig. 1B).

**Figure 1.**
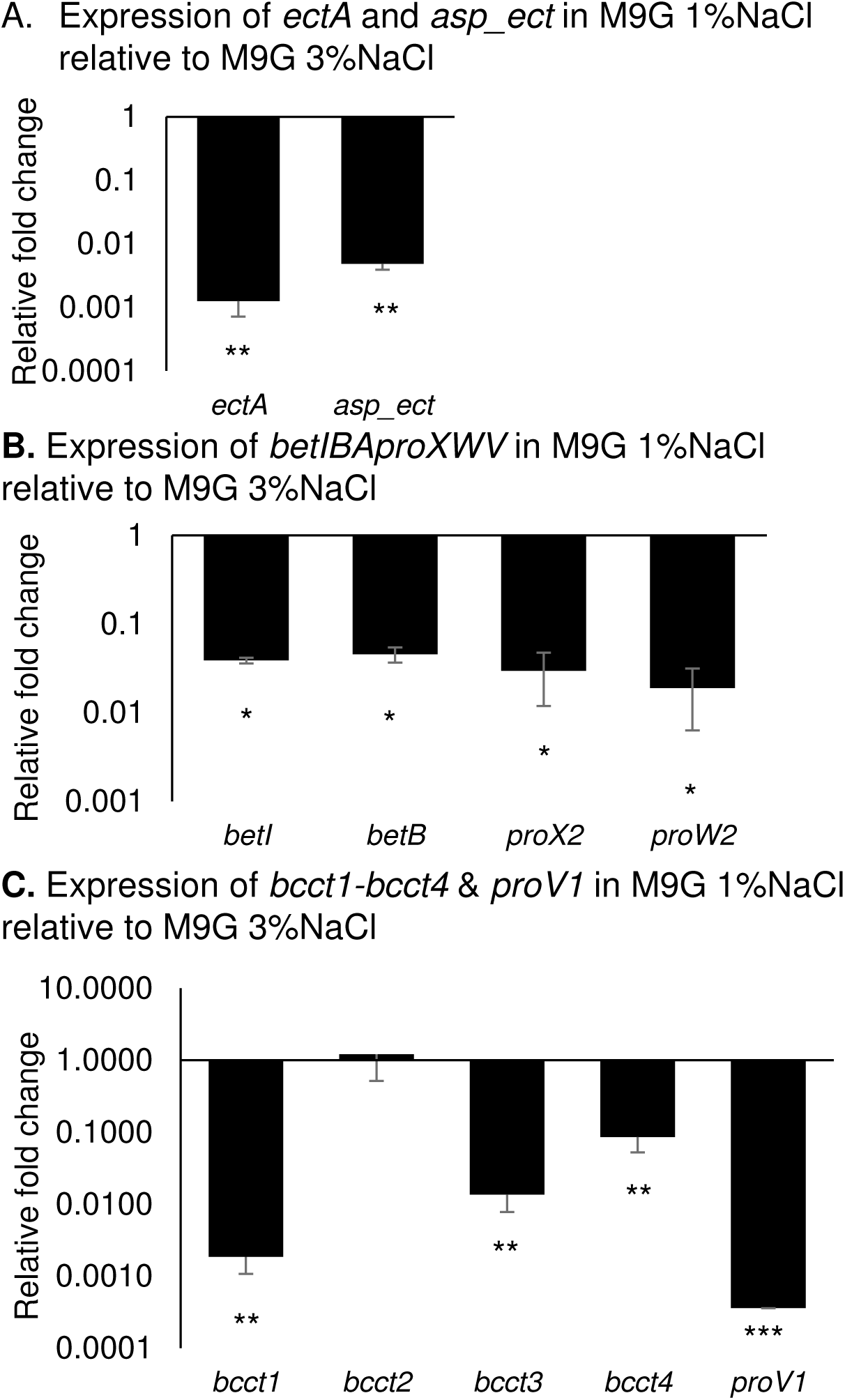
RNA was isolated from RIMD2210633 after growth in M9G1% and M9G3% at an OD_595_ of 0.45. Expression analysis of **(A)** *ectA, asp_ect*, **(B)** *betI, betB, proX2, proW2* **(C)** *bcct1, bcct2, bcct3, bcct4* and *proV1* by quantitative real time PCR (qPCR). 16S was used for normalization. Expression levels shown are levels in M9G1% relative to M9G3%. Mean and standard error of two biological replicates are shown. Statistics were calculated using a Student’s t-test (*, P < 0.05; **, P < 0.01; ***, P < 0.001).

We determined the expression levels of *bcct* genes in *V. parahaemolyticus* in both M9G1% and M9G3%. Expression of *bcct1*, *bcct3*, and *bcct4* are significantly repressed in M9G1%, 500-fold, 71.4-fold, and 11.6-fold, respectively, when compared with expression in M9G3% (Fig. 1C). The *bcct2* gene remained unchanged. We previously reported that *bcct2* is not induced by salinity (29), and our data indicates that it has a basal level of transcription in the cell based on similar Ct values in both salinities tested (data not shown). We then examined the expression pattern of the ProU1 transporter genes in *V. parahaemolyticus*. The *proV1* gene is significantly repressed in M9G1%, with a 2,786-fold change as compared to M9G3% (Fig. 1C). Overall, the data demonstrates osmoregulation of ectoine and glycine betaine biosynthesis genes and transporter genes *bcct1, bcct3, bcct4* and *proVWX*.

### CosR represses compatible solute biosynthesis and transport genes in low salinity

Next, we wanted to determine how these compatible solute systems are repressed in *V. parahaemolyticus*. Since we know CosR is a repressor of ectoine biosynthesis genes, we wondered whether it played a broader role in repression of the osmotic stress response genes, *betIBAproXWV* operon, *proVWX*, and *bcct* transporters, in low salt conditions. We examined expression of these genes in wild-type and an in-frame deletion mutant of *cosR*. RNA was isolated from the Δ*cosR* mutant strain at mid-exponential phase (OD_595_ 0.45) after growth in M9G1% and compared to wild-type RIMD2210633 grown under identical conditions. Using qPCR analysis, we determined the expression levels of *ectA* and *asp_ect* and show they are significantly induced, 818.5-fold and 308.2-fold, respectively, in a Δ*cosR* mutant compared to wild-type in M9G1% (Fig. 2A). Next, we examined expression levels of *betIBAproXWV* after growth in M9G1% in the Δ*cosR* and wild-type strains using qPCR. The *betI*, *betB*, *proX2* and *proW2* genes are significantly induced in the Δ*cosR* mutant with *betI* expressed 13.75-fold, *betB* 10.18-fold, *proX2* 8.23-fold, and *proW2* 16.38-fold more than in the wild-type strain (Fig. 2B). Similarly, we examined levels of the *bcct* genes and *proV1* in a Δ*cosR* mutant versus the wild-type in M9G1%. Relative expression levels of *bcct1* are 155.66-fold higher and levels of *bcct*3 are 34.97-fold higher than wild-type levels, while levels of *bcct2* and *bcct4* are unchanged (Fig. 2C). The *proV1* gene is induced 379.5-fold in the Δ*cosR* mutant over the wild-type strain (Fig. 2C). In sum, these data demonstrate that CosR is a repressor of *ectABCasp_ect*, *betIBAproXWV*, *bcct1, bcct3* and *proVWX* under low salinity conditions.

**Figure 2.**
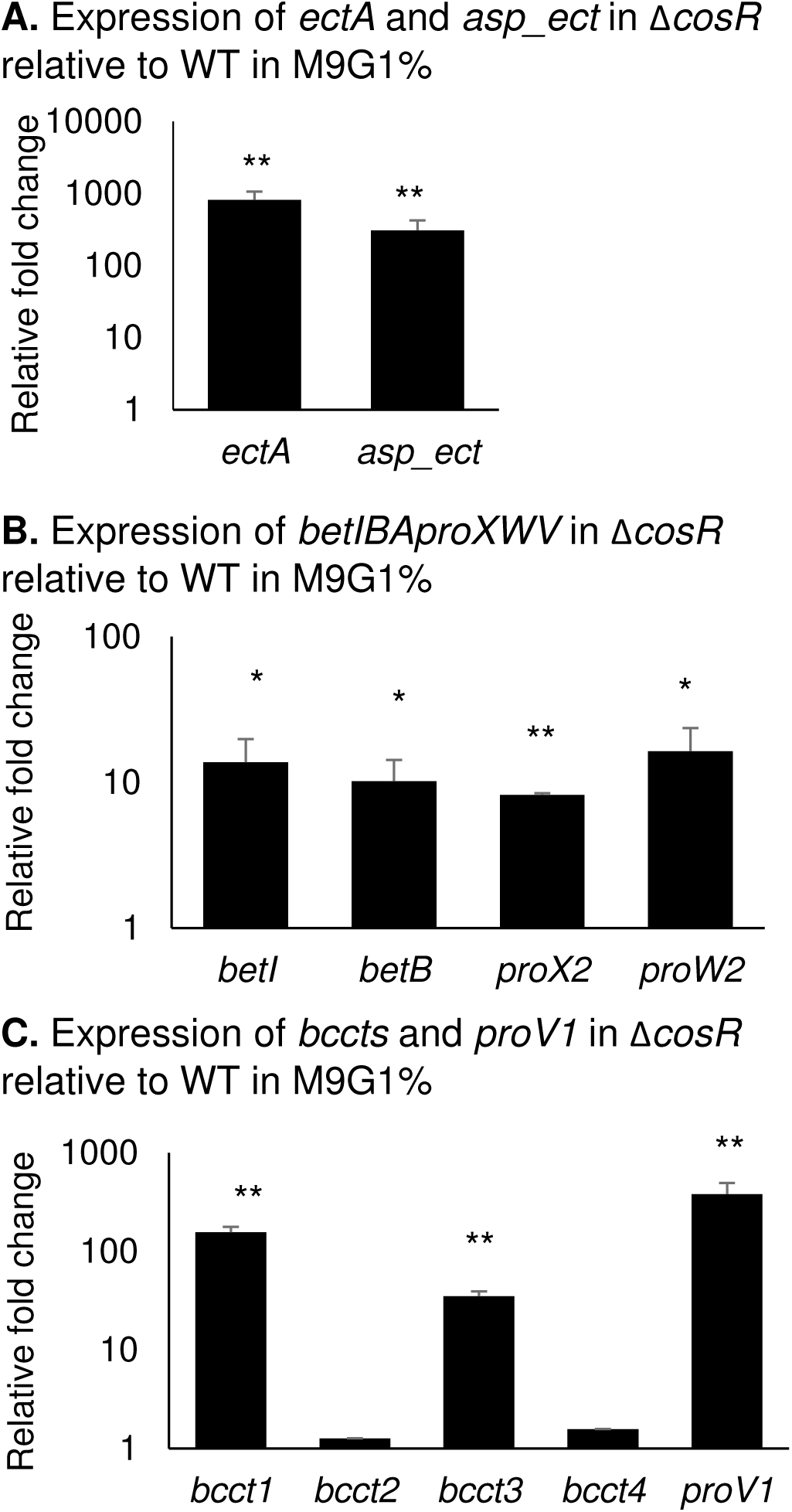
RNA was isolated from RIMD2210633 and Δ*cosR* after growth in M9G1% at an OD_595_ of 0.45. Expression analysis of **(A)** *ectA, asp_ect*,**(B)** *betI, betB, proX2, proW2* **(C)** *bcct1, bcct2, bcct3, bcct4* and *proV1* by qPCR. 16S was used for normalization. Expression levels shown are levels in Δ*cosR* relative to wild-type. Mean and standard error of two biological replicates are shown. Statistics were calculated using a Student’s t-test (*, P < 0.05; **, P < 0.01).

### CosR binds directly to the promoter of the *betIBAproXWV* operon and represses transcription

Previously, we found that CosR binds to the regulatory region of the ectoine biosynthesis operon and represses transcription (17). To determine whether CosR regulation of the glycine betaine biosynthesis operon is also direct, we performed DNA binding assays with purified CosR protein and DNA probes of the regulatory region of this operon. The regulatory region was split into five overlapping probes, P*betI* probes A-E, of sizes 125-bp, 112-bp, 142-bp, 202-bp, and 158-bp (Fig. 3A). CosR bound to probe A, which is directly upstream of the start codon for *betI*, and it also bound to probes B and D (Fig. 3B). CosR did not bind to probes C and E, demonstrating specificity of CosR binding (Fig. 3B).

**Figure 3.**
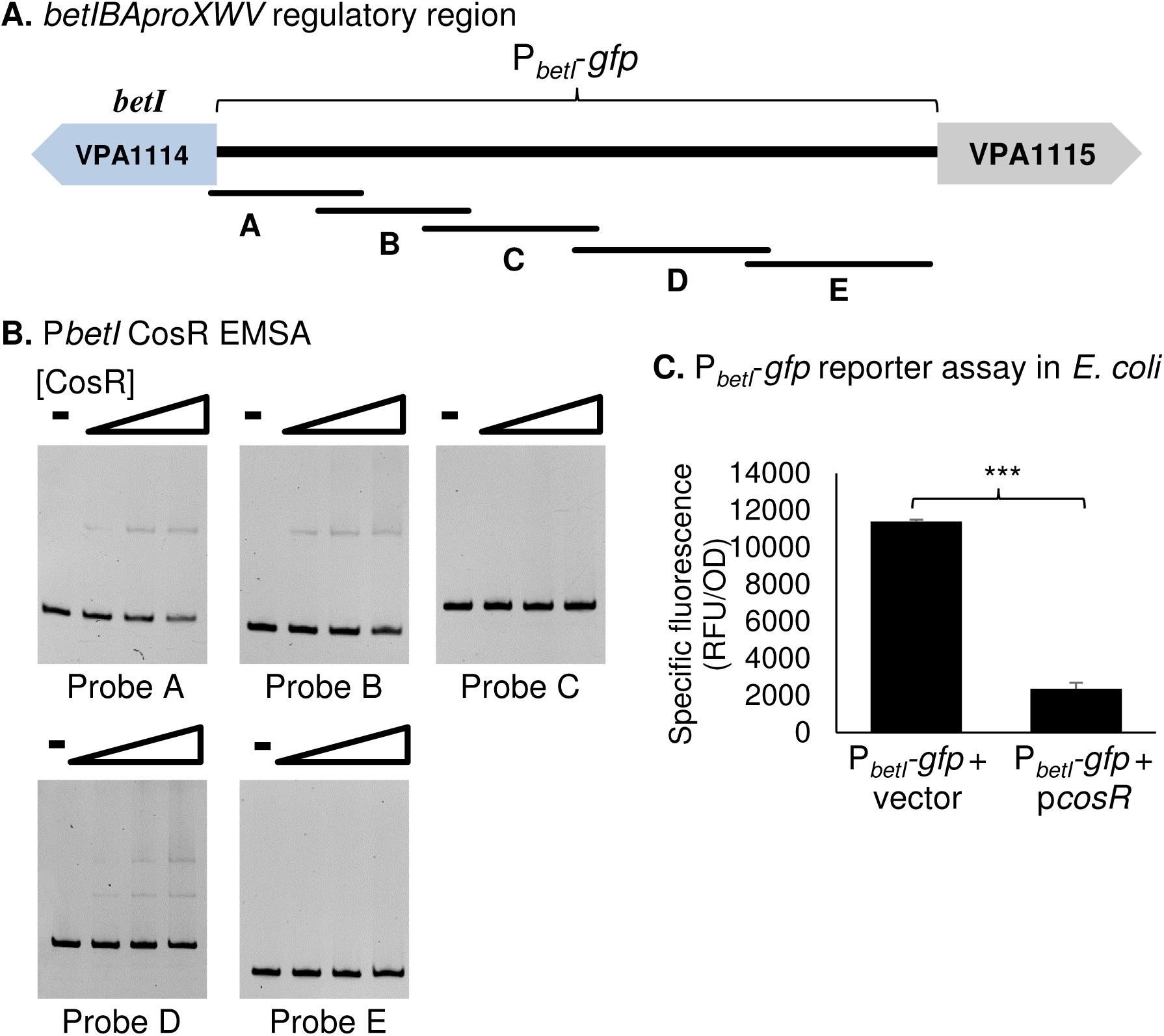
**(A)** The regulatory region of *betIBAproXWV* was divided into five probes for EMSAs, P*betI* A-E, 125-bp, 112-bp, 142-bp, 202-bp and 158-bp, respectively. The regulatory region used for the GFP reporter assay is indicated with a bracket. **(B)** An EMSA was performed with purified CosR-His (0 to 0.62 µM) and 30 ng of each P*betI* probe, with DNA:protein molar ratios of 1:0, 1:1, 1:5, and 1:10. **(C)** A P*_betI_*-*gfp* reporter assay was performed in *E. coli* strain MKH13 containing an expression plasmid with full-length *cosR* (p*cosR*). Specific fluorescence of the CosR-expressing strain was compared to a strain harboring empty expression vector. Mean and standard deviation of two biological replicates are shown. Statistics were calculated using a Student’s t-test (***, P < 0.001).

To demonstrate that direct binding by CosR results in transcriptional repression of the *betIBAproXWV* operon, we performed a GFP-reporter assay in *E. coli* strain MKH13. Full-length *cosR* was expressed from a plasmid (pBBR*cosR*) in the presence of a *gfp*-expressing reporter plasmid under the control of the glycine betaine biosynthesis system regulatory region (P*_betI_-gfp*). Relative fluorescence and OD_595_ were measured after overnight growth in M9G1%. Specific fluorescence was calculated by normalizing to OD and compared to specific fluorescence in a strain with an empty expression vector (pBBR1MCS) that also contained the P*_betI_-gfp* reporter plasmid. The activity of the P*_betI_-gfp* reporter was significantly repressed 4.84-fold as compared to the empty vector strain (Fig. 3C). This indicates that CosR directly represses transcription of the *betIBAproXWV* genes.

### CosR binds directly to the promoter of *bcct1* and *bcct3*

Next, we wanted to investigate whether CosR repression of *bcct1* and *bcct3* was direct. We designed probes upstream of the translational start for *bcct1* and *bcct3*. The 291-bp regulatory region of P*bcct1*, which includes 15-bp of *bcct1* and 276-bp of the intergenic region, was split into three overlapping probes, P*bcct1* probes A, B, and C, 120-bp, 110-bp, and 101-bp, respectively (Fig. 4A). DNA binding assays were performed with increasing concentrations of CosR. CosR bound directly to the P*bcct1* probe B (Fig. 4B) but did not bind to the other probes tested, indicating that regulation by CosR is direct and binding is specific. We then performed GFP reporter assays in *E. coli* using a GFP expression plasmid under the control of the regulatory region of *bcct1*. and a CosR expression plasmid (pBBR*cosR*). Specific fluorescence in the presence of CosR was compared to a strain with empty expression vector (pBBR1MCS). The activity of the P*_bcct1_-gfp* reporter was not significantly different than the strain harboring empty expression vector (Fig. 4C), indicating that CosR does not directly repress *bcct1*.

**Figure 4.**
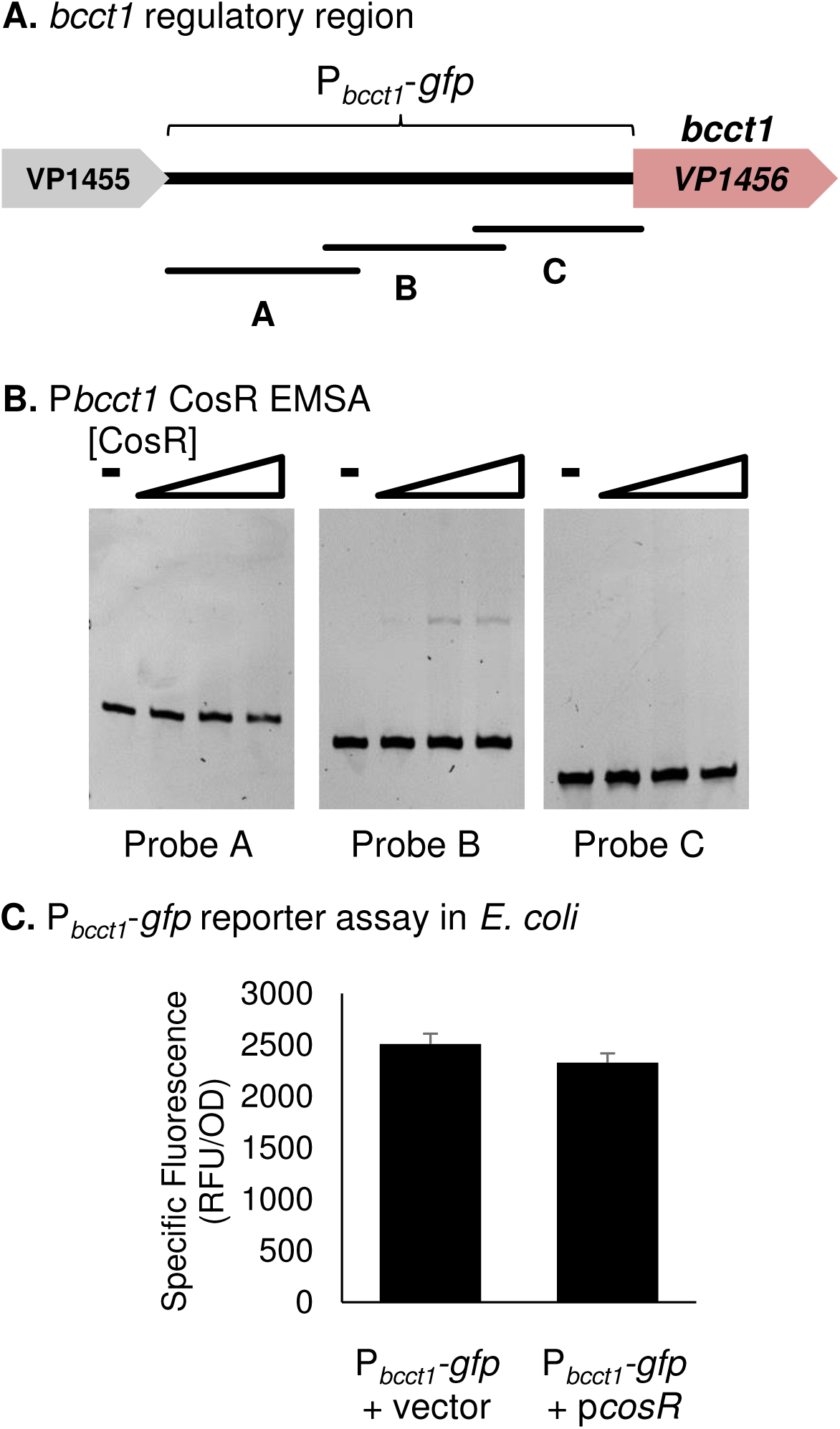
**(A)** The regulatory region of *bcct1* was divided into three similarly sized probes for EMSAs, P*bcct1* A-C, 120-bp, 110-bp, and 101-bp, respectively. The regulatory region used for the GFP reporter assay is indicated with a bracket. **(B)** An EMSA was performed with purified CosR-His (0 to 0.69 µM) and 30 ng of P*bcct1* probe with DNA:protein molar ratios of 1:0, 1:1, 1:5, and 1:10. **(C)** A P*_bcct1_*-*gfp* reporter assay was performed in *E. coli* strain MKH13 containing an expression plasmid with full-length *cosR* (p*cosR*). Specific fluorescence of the CosR-expressing strain was compared to a strain harboring empty expression vector (pBBR1MCS). Mean and standard deviation of two biological replicates are shown. Statistics were calculated using a Student’s t-test (**, P < 0.01).

Two overlapping probes designated P*bcct3* probe A and B, 108-bp and 107-bp, respectively, were designed encompassing 196-bp of the regulatory region of *bcct3* (Fig. 5A). Because *bcct3* is divergently transcribed from *cosR*, we used approximately half of the regulatory region for the P*bcct3* EMSA. An EMSA showed that CosR bound directly to the P*bcct3* probe A, which is proximal to the start of the gene, but not probe B (Fig. 5B). We then performed GFP reporter assays in *E. coli* using a GFP expression plasmid under the control of the regulatory region of *bcct3*, utilizing the entire 397-bp intergenic region between *bcct3* and *cosR*. Transcriptional activity of the P*_bcct3_-gfp* reporter is significantly repressed in a CosR-expressing strain, indicating that CosR directly represses transcription of *bcct3* (Fig. 5C). In addition, we showed that CosR does not bind to the regulatory region of *bcct2* and *bcct4* (Fig. 5D), which is in agreement with the *cosR* mutant expression data (Fig. 2C). These data suggest that *bcct2* and *bcct4* are under the control of a yet to be described regulator.

**Figure 5.**
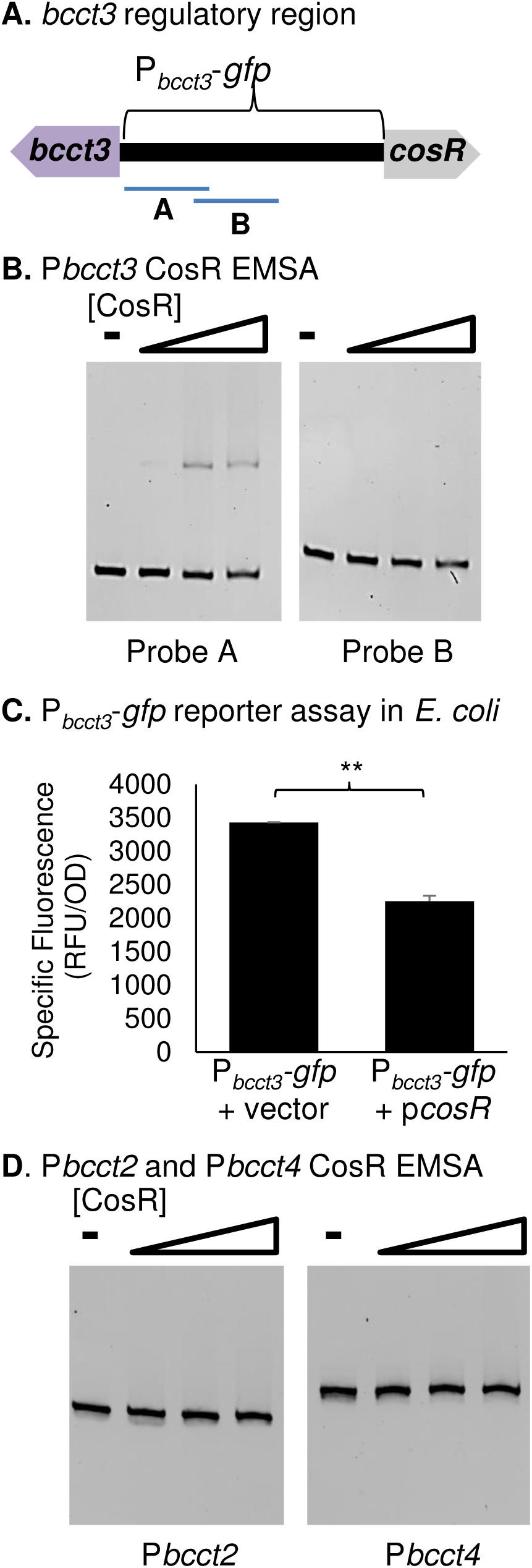
**(A)** A 196-bp portion of the regulatory region of *bcct3* was split into two probes for EMSAs, P*bcct3* A and B, 108-bp and 107-bp, respectively. The regulatory region used for the GFP reporter assay is indicated with a bracket. **(B)** An EMSA was performed with purified CosR-His (0 to 0.65 µM) and 30 ng of P*bcct3* probe with DNA:protein molar ratios of 1:0, 1:1, 1:5, and 1:10. **(C)** P*_bcct3_*-*gfp* reporter assay was performed in *E. coli* strain MKH13 containing an expression plasmid with full-length *cosR* (p*cosR*). Specific fluorescence of the CosR-expressing strain was compared to a strain harboring empty expression vector (pBBR1MCS). Mean and standard deviation of two biological replicates are shown. Statistics were calculated using a Student’s t-test (**, P < 0.01). **(D)** An EMSA was performed with CosR-His (0 to 0.18 µM) and probes of the regulatory regions of *bcct2* and *bcct4*. Each lane contains 30 ng of DNA and DNA:protein molar ratios of 1:0, 1:1, 1:5, and 1:10.

### CosR binds directly to the *proVWX* regulatory region

We also examined direct regulation of the *proVWX* operon on chromosome 1 by CosR. The regulatory region upstream of the *proV1* gene was divided into four probes, 160-bp, 134-bp, 108-bp and 109-bp (Fig. 6A). A DNA binding assay was performed with increasing concentrations of CosR and 30 ng of each probe. A shift in the DNA bands of probe D, which is proximal to the start codon of *proV1*, indicates that CosR binds directly to this region (Fig. 6B). CosR did not bind to the other probes tested, indicating that CosR binding is specific.

**Figure 6.**
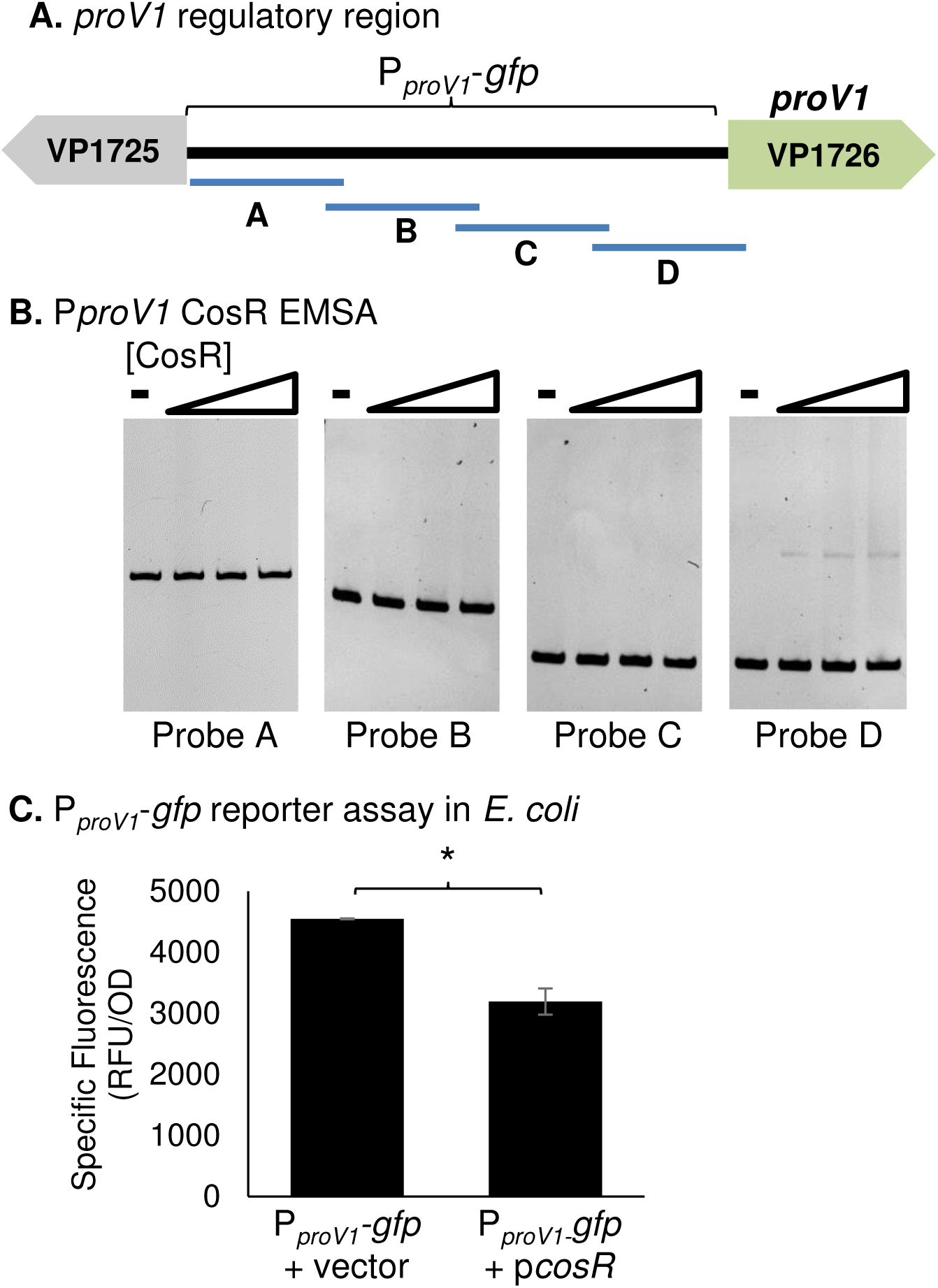
**(A)** The 447-bp regulatory region of the *proV1* gene was divided into four probes for EMSAs, P*proV1* A-D, 160-bp, 134-bp, 108-bp and 109-bp, respectively. The regulatory region used for the GFP reporter assay is indicated with a bracket. **(B)** An EMSA was performed with purified CosR-His (0 to 0.64 µM) and 30 ng of each P*proV1* probe with DNA:protein molar ratios of 1:0, 1:1, 1:5, and 1:10. **(C)** A reporter assay was conducted in *E. coli* MKH13 harboring the P*_proV1_*-gfp reporter plasmid and the expression plasmid p*cosR*. Specific fluorescence of the CosR-expressing strain was compared to an empty vector strain. Mean and standard deviation of two biological replicates are shown. Statistics were calculated using a Student’s t-test (*, P < 0.05).

We also performed a GFP-reporter assay in *E. coli* utilizing the *cosR* expression plasmid (pBBRcosR) and a GFP reporter plasmid under the control of the *proVWX* regulatory region (P*_proV1_-gfp*). We found that in a CosR-expressing strain, expression of the P*_proV1_-gfp* reporter was significantly repressed when compared to an empty expression vector strain (Fig. 6C). This indicates that CosR is a direct repressor of the *proVWX* operon.

### CosR is a MarR-type regulator that does not participate in an autoregulatory feedback loop

In *V. cholerae*, expression levels of *cosR* are upregulated in 0.5 M NaCl as compared to levels in 0.2 M NaCl (18). It was suggested that one reason for the upregulation of *cosR* in higher salinity could be that it is involved in an autoregulatory feedback loop (18). In *V. parahaemolyticus*, we found that levels of *cosR* are not significantly upregulated in moderate salinity (3% NaCl) as compared to low salinity (1% NaCl) (data not shown). We have already shown that CosR binds to the intergenic region between *bcct3* and *cosR*, but the binding site location is proximal to the start codon of *bcct3*, more than 300 bp upstream of the *cosR* gene (Fig. 5A & B). Therefore, to investigate CosR autoregulation, we designed two probes, 105-bp and 142-bp, which comprise a 220-bp portion of the regulatory region upstream of *cosR* (VP1906) (Fig. 7A) and used this in a DNA binding assay with various concentrations of purified CosR (Fig. 7B). There are no shifts observed in the binding assay, indicating that CosR does not bind (Fig. 7B). We then performed a GFP reporter assay in *E. coli*, utilizing the entire 397-bp intergenic region between *bcct3* and *cosR*, to determine if CosR directly represses transcription of its own gene. The transcriptional activity of P*_cosR_-gfp* in the presence of CosR was not significantly different from the empty-vector strain (Fig. 7C). We therefore conclude that under these conditions, in *V. parahaemolyticus* CosR does not autoregulate, and that the CosR binding site proximal to the *bcct3* gene does not affect transcription of the *cosR* gene.

**Figure 7.**
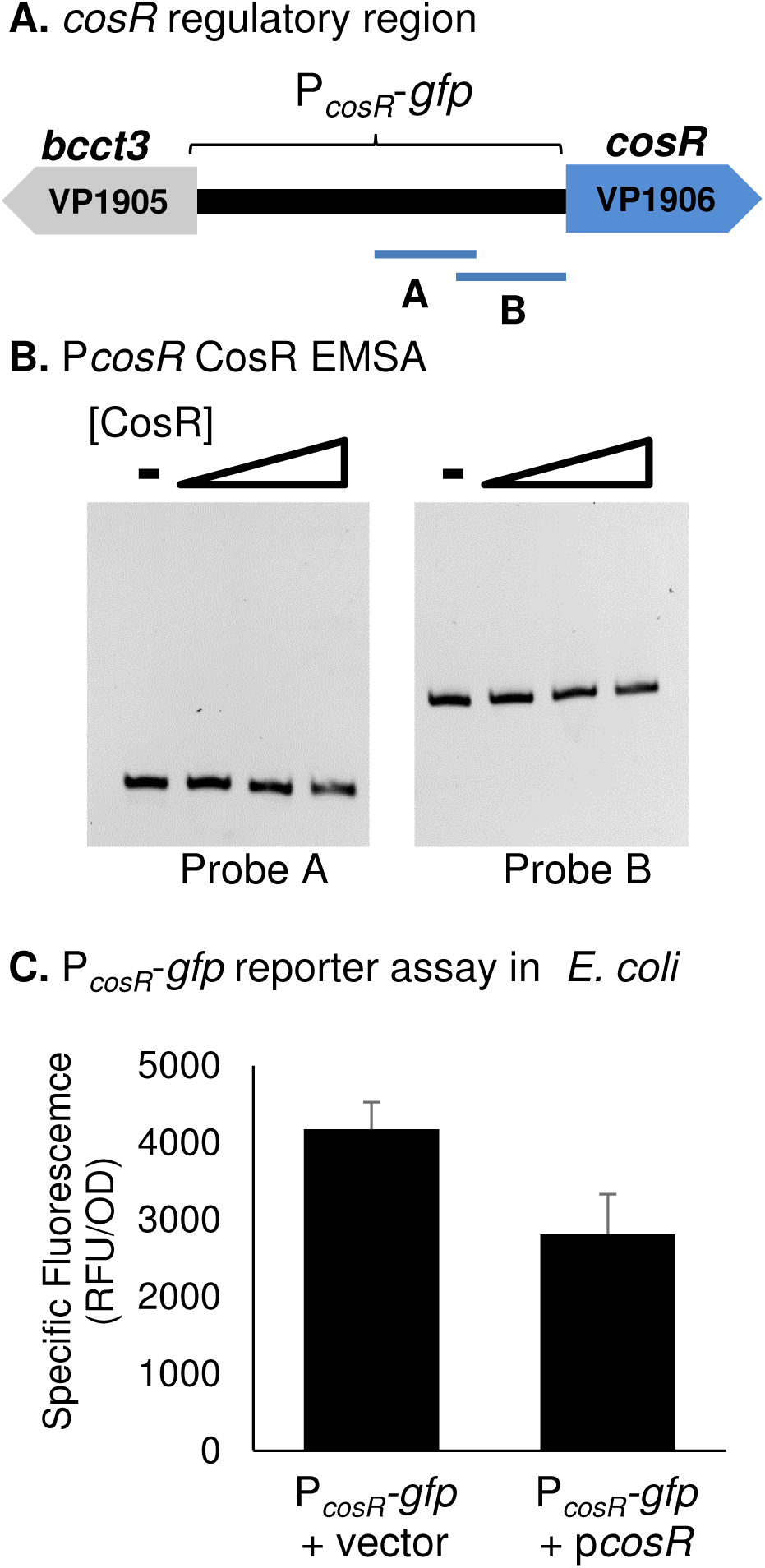
**(A)** A 220-bp section of the regulatory region of *cosR* was split into two similarly sized probes for EMSAs, P*cosR* A and B, 105-bp and 142-bp, respectively. The regulatory region used for the GFP reporter assay is indicated with a bracket. **(B)** An EMSA was performed with increasing concentrations of purified CosR-His (0 to 0.66 µM) and 30 ng of each probe with DNA:protein molar ratios of 1:0, 1:1, 1:5, and 1:10. **(C)** A P*_cosR_*-*gfp* reporter assay was performed in *E. coli* strain MKH13 the p*cosR* expression plasmid. Specific fluorescence of the CosR-expressing strain was compared to a strain harboring empty expression vector. Mean and standard deviation of two biological replicates are shown.

### BetI represses its own operon in the absence of choline

Previously, it was shown that BetI represses its own operon in several bacterial species and this repression is relieved in the presence of choline (27, 31, 32). To demonstrate BetI regulates its own operon in *V. parahaemolyticus*, we performed a plasmid-based GFP reporter assay utilizing the P*_betI_-gfp* reporter in RIMD2210633 strain and a Δ*betI* mutant strain. Strains were grown overnight in M9G3%, with and without choline, and specific fluorescence was calculated. Expression of the reporter is significantly induced in the Δ*betI* mutant when no choline is present, indicating that BetI is a negative regulator of its own operon (Fig. 8A). In the presence of choline, there is no longer a significant difference in reporter activity between the wild-type strain and the Δ*betI* mutant strain, indicating that repression by BetI is relieved (Fig. 8B).

**Figure 8.**
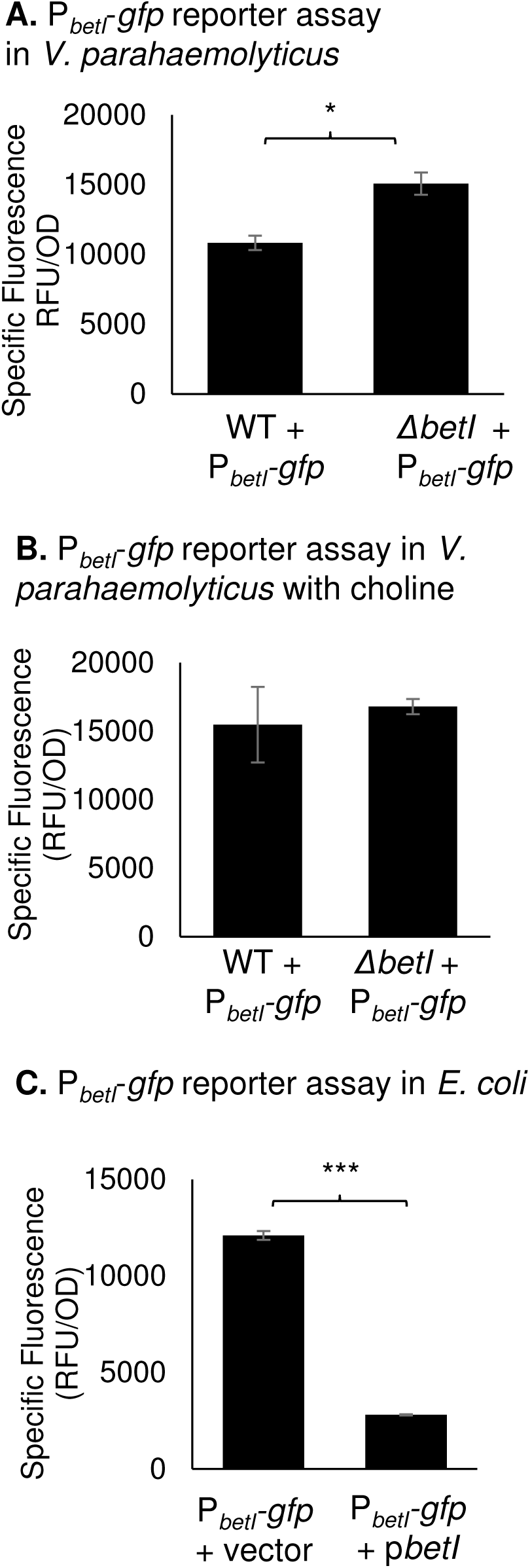
**(A)** Expression of a P*_betI_*-*gfp* transcriptional fusion reporter in wild-type and a Δ*betI* mutant. Relative fluorescence intensity (RFU) and OD_595_ were measured after growth in **(A)** M9G3% or **(B)** M9G3% with the addition of choline. Specific fluorescence was calculated by dividing RFU by OD. Mean and standard deviation of two biological replicates are shown. Statistics were calculated using a Student’s t-test (*, P < 0.05). **(C)** A reporter assay was conducted in *E. coli* MKH13 using the P*_betI_*-*gfp* reporter plasmid and an expression plasmid with full-length *betI* (p*betI*). The specific fluorescence was calculated and compared to a strain with an empty expression vector (pBBR1MCS). Mean and standard deviation of two biological replicates are shown. Statistics were calculated using a Student’s t-test (***, P < 0.001).

To determine whether regulation of *betIBAproXWV* by BetI is direct, we performed a GFP reporter assay in *E. coli* MKH13 strain. The P*_betI_-gfp* reporter utilized in our *in vivo* reporter assay was introduced into the *E. coli* MKH13 strain (which lacks its own *betIBA* operon) along with an expression vector harboring full-length *betI* under the control of an IPTG-inducible promoter. In the BetI-expressing strain, P*_betI_-gfp* expression was significantly repressed, indicating that BetI is a direct repressor of its own operon in *V. parahaemolyticus* (Fig. 8C).

### The LuxR homolog OpaR is a positive regulator of *betIBAproXWV* in *V. parahaemolyticus*

It was demonstrated in *V. harveyi* that LuxR, the quorum sensing master regulator, induces *betIBAproXWV* expression and that this regulation is direct (32). We examined expression of the P*_betI_-gfp* reporter in wild-type and the Δ*opaR* mutant in *V. parahaemolyticus*. Expression of the reporter is significantly repressed in Δ*opaR*, indicating that OpaR is a positive regulator of the glycine betaine biosynthesis operon in *V. parahaemolyticus* (Fig. 9A). We also examined whether regulation of P*betI* by OpaR was direct utilizing an EMSA with purified OpaR protein. The P*betI* probes A-E used previously in the CosR EMSA (Fig. 3A) were incubated with purified OpaR. OpaR bound to all P*betI* probes, indicating that regulation of *betIBAproXWV* by OpaR is direct (Fig. 9B).

**Figure 9.**
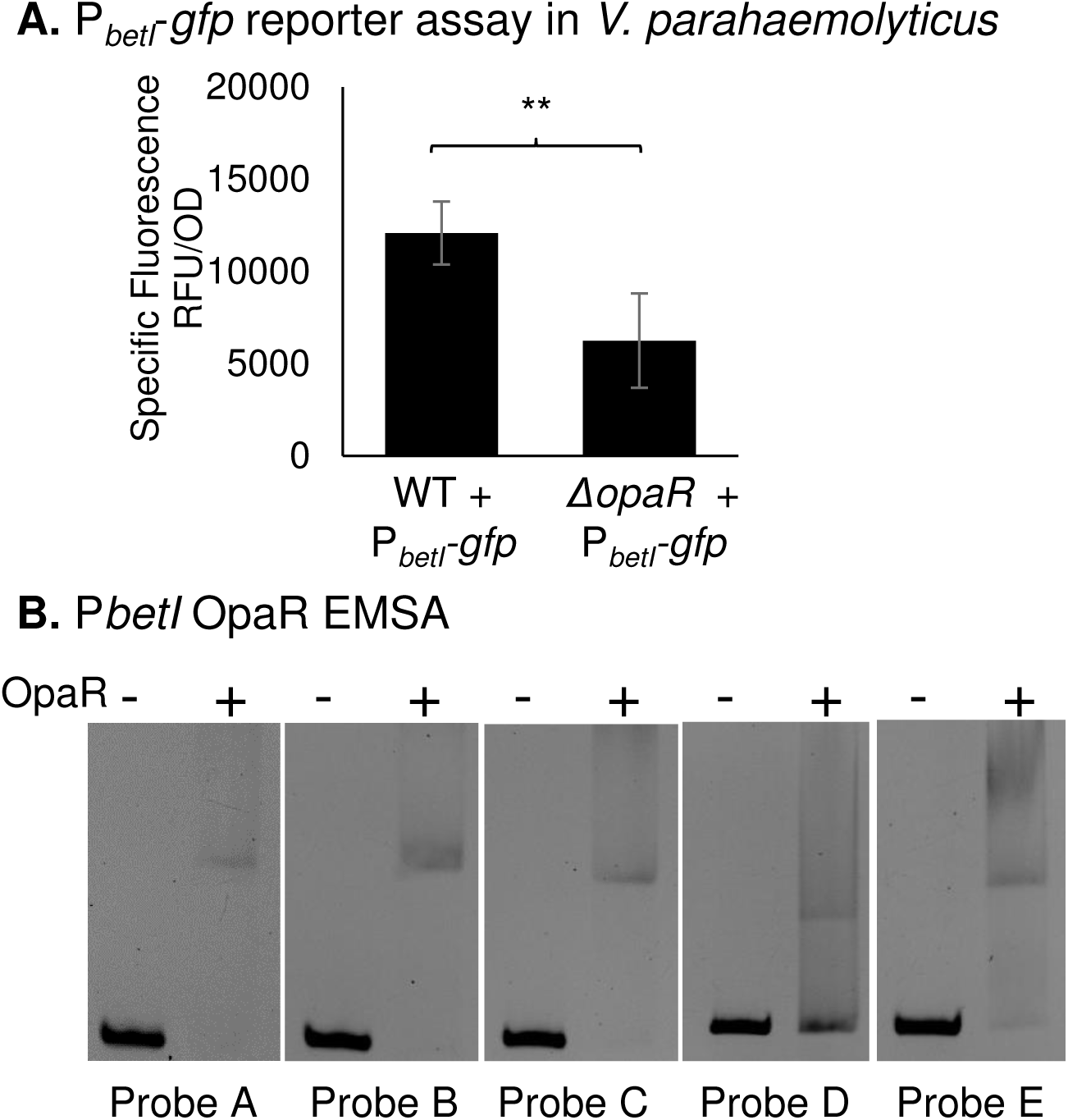
**(A)** Expression of a P*_betI_*-*gfp* transcriptional fusion reporter in wild-type and Δ*opaR* mutant strains. Relative fluorescence intensity (RFU) and OD_595_ were measured after growth in M9G3%. Specific fluorescence was calculated by dividing RFU by OD. Mean and standard deviation of two biological replicates are shown. Statistics were calculated using a one-way ANOVA with a Tukey-Kramer *post hoc* test (**, P < 0.01). **(B)** An EMSA was performed with 30 ng of each P*betI* probe A-E utilized previously in the CosR EMSA and purified OpaR protein (between 0.47 and 0.82 µM) in a 1:20 molar ratio of DNA:protein.

### Distribution of compatible solute biosynthesis and transport systems in *Vibrionaceae*

CosR, a MarR family regulator, in *V. parahaemolyticus* is a 158 amino acid protein that is divergently transcribed from *bcct3* on chromosome 1. Our BLAST analysis showed that a CosR homolog is present in over 50 *Vibrio* species and in all cases the *cosR* homolog was divergently transcribed from a *bcct* transporter. Within these *Vibrio* species, homology ranged from 98% to 73% amino acid identity. We found that in *V. splendidus, V. crassostreae, V. cyclitrophicus, V. celticus*, *V. lentus* and *Aliivibrio wodanis*, the CosR homolog is present directly downstream of the *betIBAproXWV* operon on chromosome 2 (**Fig. 10**). CosR in these species share ∼73-75% amino acid identity with CosR in *V. parahaemolyticus*. In *V. tasmaniensis* strains and *Vibrio sp.* MED222, the CosR homolog is also downstream of the betaine biosynthesis operon and the operon for ectoine biosynthesis clusters in the same genome location (**Fig. 10**). In two *Aliivibrio wodanis* strains, AWOD1 and 06/90/160, *cosR* homologs were clustered with putative transporters and the glycine betaine biosynthesis operon. In all strains of *Aliivibrio fischeri*, the *cosR* homolog (which shares 73% amino acid identity with CosR from *V. parahaemolyticus*) clusters with two uncharacterized transporters. However, a second MarR family regulator, a 141 amino acid protein, which we name *ectR*, clusters with the ectoine biosynthesis genes in this species. EctR shares only 31% identity with less than 60% query coverage to CosR from *V. parahaemolyticus* and a similar level of low amino acid identity to EctR1 from *Methylmicrobium alcaliphilum*. EctR was also clustered with the *ectABCasp_ect* genes in all strains of *Aliivibrio finisterrensis, Aliivibrio sifiae*, and most *A. wodanis* strains. Thus, in *Aliivibrio* species, it appears that the ectoine gene cluster has a new uncharacterized regulator of the MarR family, which was confined to this group.

**Figure 10.**
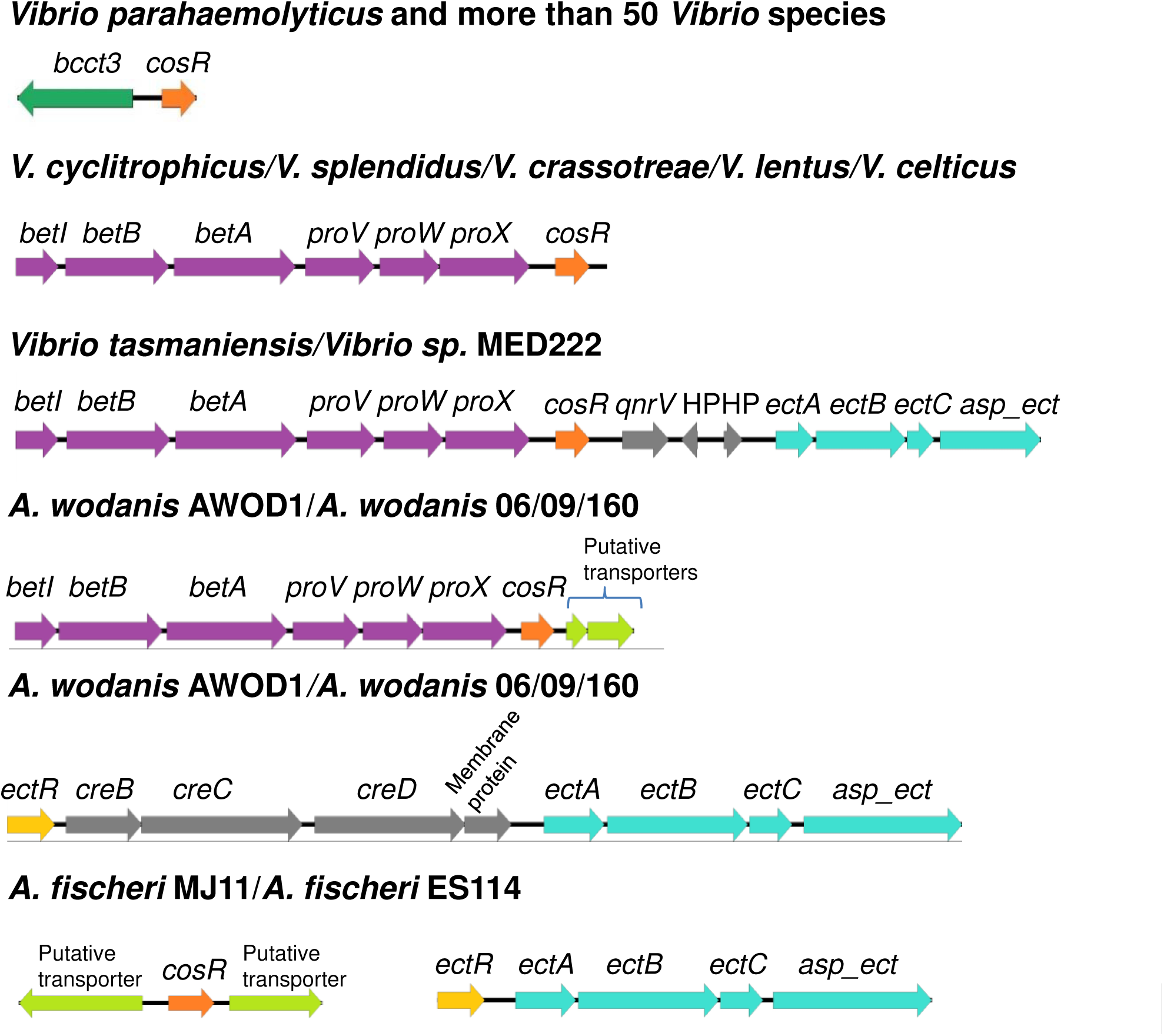
Schematic of the genomic context of CosR homologs from select Vibrionaceae species. Open reading frames are designated by arrows.

## Discussion

Here we have shown that the compatible solute biosynthesis and transport genes are downregulated in *V. parahaemolyticus* in low salinity. Our genetic analysis, binding analysis, and reporter assays demonstrate that the transcriptional regulator CosR is a direct repressor of *betIBAproXWV, bcct3*, and *proVWX* in low salinity. Additionally, we show that under the conditions tested, CosR is not autoregulated in *V. parahaemolyticus*. Our bioinformatics analysis indicates that CosR repression of compatible solute systems is likely widespread within the *Vibrio* genus.

Although CosR binds directly to the regulatory region of *bcct1*, transcription was not directly repressed in our reporter assay. Based on our expression data combined with our DNA-binding assays, we speculate it is probable that CosR also directly represses *bcct1* expression, but we could not detect significant differences between the CosR- and empty vector-expressing strains due to the low level of activation of the *bcct1* regulatory region in *E. coli*.

CosR characterized from *Vibrio* species show ∼50% amino acid identity to EctR1, a MarR-type regulator in the halotolerant methanotroph *Methylmicrobium alcaliphilum* (49). In this species, *ectR1* is divergently transcribed from the same promoter as *ectABC-asp_ect*. Mustakhimov and colleagues showed that EctR1 repressed expression of the *ectABC-ask* operon in response to low salinity (49). Purified EctR1 bound specifically to the promoter of *ectABC-ask*, indicating direct regulation by EctR1 (49). EctR repression of the ectoine biosynthesis genes was also shown in both *Methylophaga alcalica* and *Methylophaga thalassica*, two moderately halophilic methylotrophs (50, 51). In *V. cholerae*, CosR was also identified as a repressor of ectoine biosynthesis genes though it does not cluster with *ectABC-asp_ect* (18). The *cosR* gene in *V. cholerae* is divergently transcribed from the *opuD* gene (a *bcct3* homolog), which is also repressed by CosR (18). Similarly, in *V. parahaemolyticus*, the *cosR* (VP1906) homolog is divergently transcribed from *bcct3* (VP1905). In this species, we demonstrated previously that CosR is a direct negative regulator of *ectABCasp_ect* and show here that it directly represses *bcct3* (17). Our bioinformatics analysis found that the CosR homolog is divergently transcribed from *bcct3* in over 50 *Vibrio* species demonstrating conservation of genomic context suggesting functional conservation. In several *Vibrio* species the CosR homolog was clustered with the *betIBAproXWV* operon, which is further suggestive of its role in regulation of compatible solute biosynthesis among *Vibrio* species. Incidentally, in *V. tasmaniensis* LGP32 (formerly *V. splendidus* LGP32) and *Vibrio* MED222, the ectoine gene cluster was present in the same genome region as the *betIBAproXWV*-*cosR* cluster.

CosR and EctR1 are members of the MarR family of transcriptional regulators, first characterized in *E. coli*, which are important regulators of a number of cellular responses, typically responding to a change in the external environment (52–54). The literature suggests that MarR-type regulators form dimers and bind to a 20-45 bp pseudo-palindromic site in the intergenic region of genes it controls (52, 55–57). The activity of MarR-type regulators can be modulated by the presence of a chemical signal, either a ligand, metal ion, or reactive oxygen species. Binding of these signals causes the protein to undergo a conformational change, thereby affecting DNA binding capability (52, 58, 59). We modeled a CosR homodimer using SWISS-MODEL and did not identify a ligand binding pocket. In *V. cholerae*, CosR activity is not affected by the presence of exogenous compatible solutes including ectoine, glycine betaine and proline, and *opuD* (*bcct* homolog) transcripts were unchanged in a *cosR* mutant. Hence, the environmental or cellular signals that modulate the activity of CosR remain unknown, as was noted by Czech and colleagues (60). Interestingly, our modelling of the EctR regulator identified in *Aliivibrio* species indicated it also does not have a ligand-binding pocket.

Autoregulation was shown for several MarR family regulators, including *ectR1* in *M. alcaliphilum* (49, 52). It was suggested previously that CosR maybe involved in an autoregulatory feedback loop in *V. cholerae* (18). In *V. parahaemolyticus* we show CosR does not bind to its own regulatory region, and our reporter assay suggests that CosR does not autoregulate. It is interesting to note that EctR1 participates in an autoregulatory feedback loop in *M. alcaliphilum* but not in *M. thalassica* (51, 61).

Ectoine biosynthesis is present in all halophilic *Vibrio* species and is essential for growth in high salt in the absence of compatible solute uptake (13). However, compatible solutes are not required under low salinity conditions. The physiological role of CosR repression of compatible solute biosynthesis in low salinity is likely to protect levels of key intracellular metabolites such as glutamate, acetyl-CoA, and oxaloacetate, all of which are affected by ectoine biosynthesis (62, 63).

Similar to ectoine biosynthesis gene expression, few direct regulators of glycine betaine biosynthesis genes have been identified. In *E. coli*, expression of *betIBA* was repressed by BetI and repression was relieved in the presence of choline (27). BetI was shown to directly regulate transcription at this locus via DNA binding assays (28). ArcA was shown to repress the *bet* operon under anaerobic conditions in *E. coli*, although direct binding was not shown (27). In *Vibrio harveyi*, it was shown that *betIBAproXWV* were repressed 2- to 3-fold when *betI* was overexpressed from a plasmid. Purified BetI bound directly to the regulatory region of the *betIBAproXWV* operon in DNA binding assays (32, 33). In these studies, it was also shown that the quorum sensing response regulator LuxR, along with the global regulator IHF, activated expression of *betIBAproXWV* (32, 33). Here we have shown that BetI represses its own operon in *V. parahaemolyticus*, as expected, and we identified a novel regulator of glycine betaine biosynthesis genes, CosR, which directly represses under low salinity conditions. We also confirm that, similar to *V. harveyi*, the quorum sensing master regulator OpaR induced *betIBAproXWV* expression in *V. parahaemolyticus* and this regulation is direct.

Biosynthesis of compatible solutes is an energetically costly process for bacteria (35). *V. parahaemolyticus* does not accumulate compatible solutes in low salinity (12, 13, 29), and therefore the transcription of biosynthesis and transport genes is unnecessary. CosR represses the genes involved in the osmotic stress response in *V. parahaemolyticus* in low salinity conditions. The high conservation of the CosR protein across *Vibrio* species and its genomic context indicates that regulation by CosR of compatible solute systems is widespread in bacteria.

## Materials and Methods

### Bacterial strains, media and culture conditions

Listed in Table 1 are all strains and plasmids used in this study. A previously described streptomycin-resistant clinical isolate of *V. parahaemolyticus*, RIMD2210633, was used as the wild-type strain (64)Makino et al., 2003). *V. parahaemolyticus* strains were grown in either lysogeny broth (LB) (Fisher Scientific, Fair Lawn, NJ) supplemented with 3% NaCl (wt/vol) (LBS) or in M9 minimal medium (47.8 mM Na_2_HPO_4_, 22 mM KH_2_PO_4_, 18.7 mM NH_4_Cl, 8.6 mM NaCl) (Sigma-Aldrich, USA) supplemented with 2 mM MgSO_4_, 0.1 mM CaCl_2_, 20 mM glucose as the sole carbon source (M9G) and 1% or 3% NaCl (wt/vol) (M9G1%, M9G3%). *E. coli* strains were grown in LB supplemented with 1% NaCl (wt/vol) or M9G1% where indicated. *E. coli* β2155, a diaminopimelic acid (DAP) auxotroph, was supplemented with 0.3 mM DAP and grown in LB 1% NaCl. All strains were grown at 37°C with aeration. Antibiotics were used at the following concentrations (wt/vol) as necessary: ampicillin (Amp), 50 µg/ml; chloramphenicol (Cm), 12.5 µg/ml; tetracycline (Tet), 1 µg/mL; and streptomycin (Str), 200 µg/ml. Choline was added to media at a final concentration of 1 mM, when indicated.

**Table 1.**
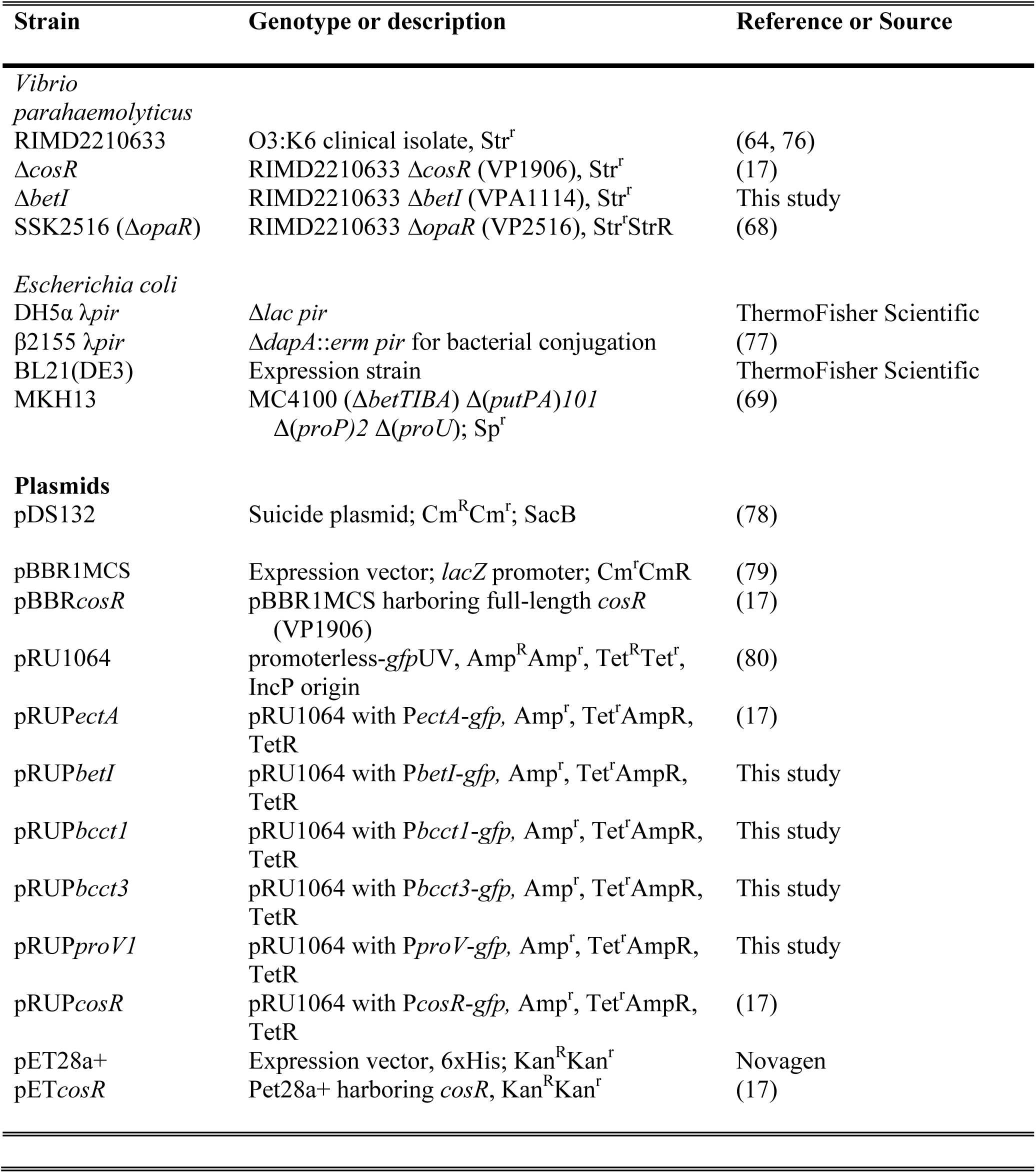
Strains and plasmids used in this study.

### Construction of the *betI* deletion mutant

An in-frame *betI* (VPA1114) deletion mutant was constructed as described previously (17). Briefly, the Gibson assembly protocol, using NEBuilder HiFi DNA Assembly Master Mix (New England Biolabs, Ipswich, MA), followed by allelic exchange, was used to generate an in-frame 63-bp truncated, non-functional *betI* gene (65, 66). Two fragments, AB and CD, were amplified from the RIMD2210633 genome using primers listed in Table 2. These were ligated with pDS132, which had been digested with SphI, via Gibson assembly to produce suicide vector pDS132 with a truncated *betI* allele (pDSΔ*betI*). pDSΔ*betI* was transformed into *E. coli* strain β2155 λ*pir*, followed by conjugation with *V. parahaemolyticus*. The suicide vector pDS132 must be incorporated into the *V. parahaemolyticus* genome via homologous recombination, as *V. parahaemolyticus* lacks the *pir* gene required for replication of the vector. Growth without chloramphenicol induces a second recombination event which leaves behind either the truncated mutant allele or the wild-type allele. Colonies were plated on sucrose for selection, as pDS132 harbors a *sacB* gene, which makes sucrose toxic to cells still carrying the plasmid. Healthy colonies were screened via PCR and sequenced to confirm an in-frame deletion of the *betI* gene.

**Table 2.**
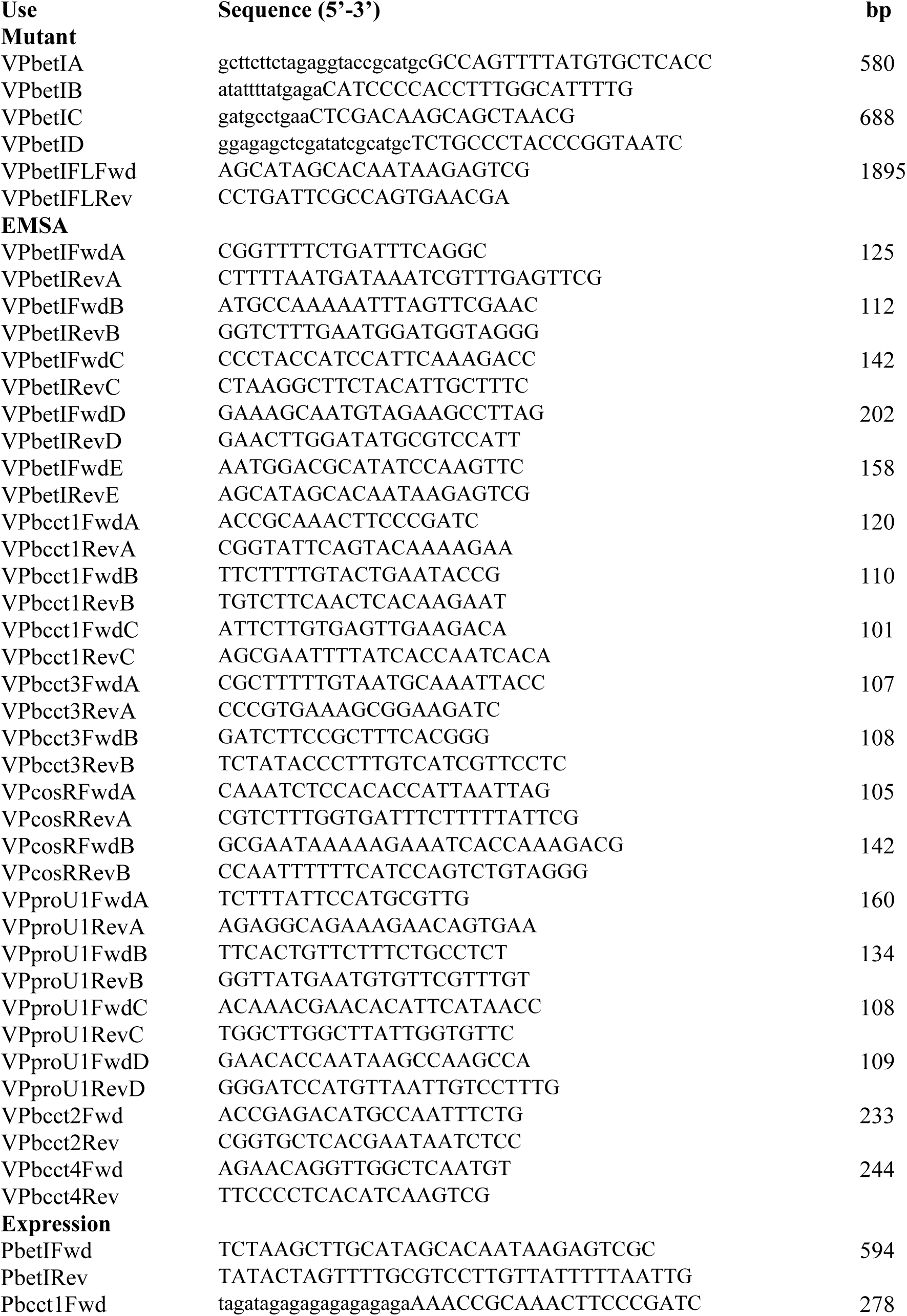
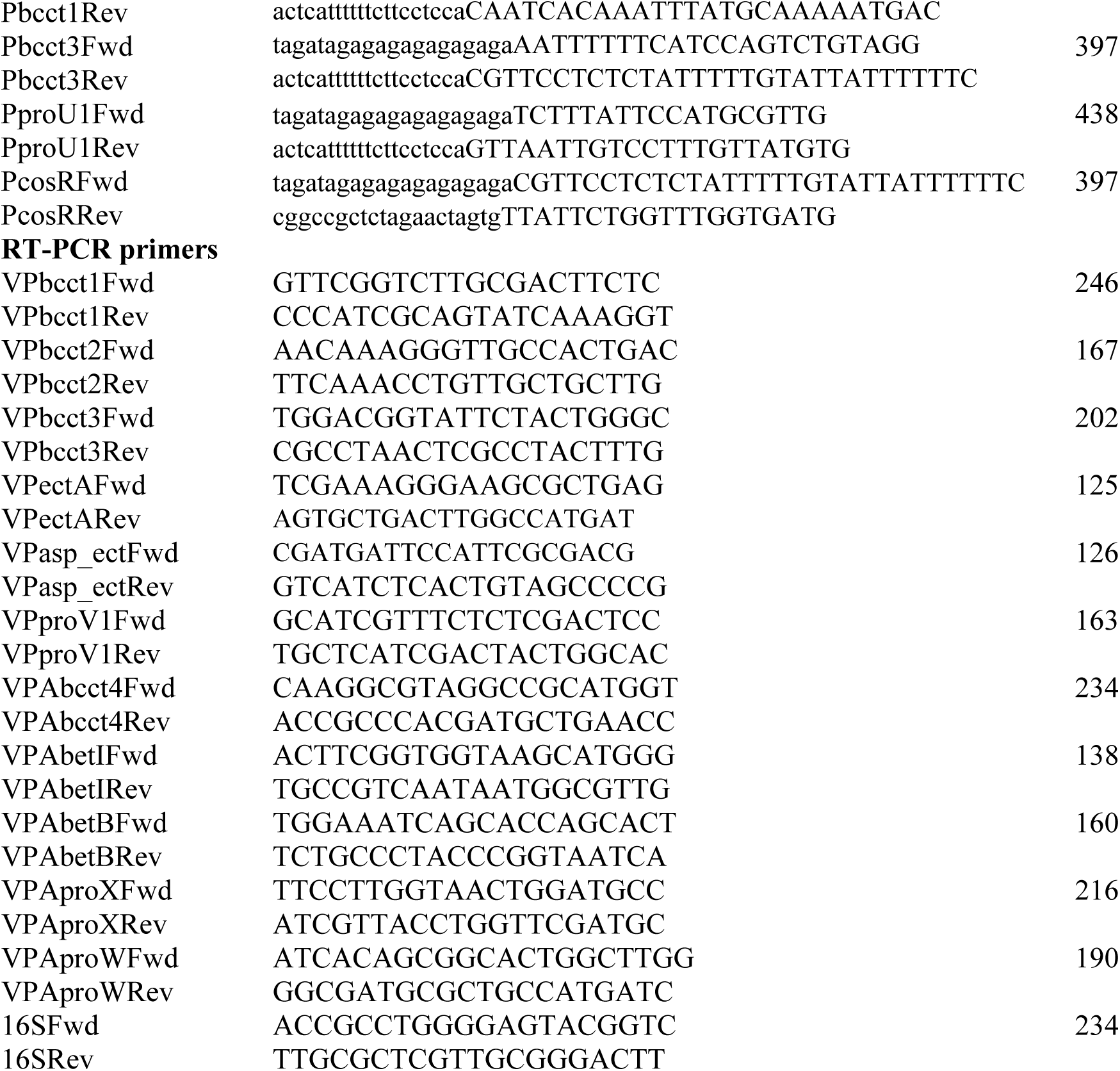
Primers used in this study.

### RNA isolation and qPCR

*Vibrio parahaemolyticus* RIMD2210633 and Δ*cosR* were grown with aeration at 37 °C overnight in LBS. Cells were pelleted, washed twice with 1X PBS, diluted 1:50 into M9G3% or M9G1% and grown with aeration to mid-exponential phase (OD_595_ 0.45). RNA was extracted from 1 mL of culture using Trizol, following the manufacturer’s protocol (Invitrogen, Carlsbad, CA). The samples were treated with Turbo DNase (Invitrogen), followed by heat inactivation of the enzyme as per manufacturer’s protocol. Final RNA concentration was quantified using a Nanodrop spectrophotometer (Thermo Scientific, Waltham, MA). 500 ng of RNA were used for cDNA synthesis by priming with random hexamers using SSIV reverse transcriptase (Invitrogen). Synthesized cDNA was diluted 1:25 and used for quantitative real-time PCR (qPCR). qPCR experiments were performed using PowerUp SYBR master mix (Life Technologies, Carlsbad, CA) on an Applied Biosystems QuantStudio6 fast real-time PCR system (Applied Biosystems, Foster City, CA). Reactions were set up with the following primer pairs listed in Table 2: VPbcct1Fwd/Rev, VPbcct2Fwd/Rev, VPbcct3Fwd/Rev, VPbcct4Fwd/Rev, VPectAFwd/Rev, VPasp_ectFwd/Rev, VPproV1Fwd/Rev, VPAbetIFwd/Rev, VPAbetBFwd/Rev, VPAproXFwd/Rev, VPAproWFwd/Rev, and 16SFwd/Rev for normalization. Expression levels were quantified using cycle threshold (CT) and were normalized to 16S rRNA. Differences in gene expression were determined using the ΔΔCT method (67).

### Protein purification of CosR

CosR was purified as described previously (17). Briefly, full-length *cosR* (VP1906) was cloned into the protein expression vector pET28a (+) containing an IPTG-inducible promoter and a C-terminal 6x-His tag (Novagen). Expression of CosR-His was then induced in *E. coli* BL21 (DE3) with 0.5 mM IPTG at OD_595_ of 0.4 and grown overnight at room temperature. Cells were harvested, resuspended in lysis buffer (50 mM NaPO4, 200 mM NaCl, 20 mM imidazole buffer [pH 7.4]) and lysed using a microfluidizer. CosR-His was bound to a Ni-NTA column and eluted with 50 mM NaPO4, 200 mM NaCl, 500 mM imidazole buffer [pH 7.4] after a series of washes to remove loosely bound protein. Protein purity was determined via SDS-PAGE. OpaR was purified as described previously (68).

### Electrophoretic Mobility Shift Assay

Five overlapping DNA fragments, designated P*betI* probe A (125-bp), probe B (112-bp), probe C (142-bp), probe D (202-bp) and probe E (158-bp), were generated from the *betIBAproXWV* regulatory region (includes 36-bp of the coding region and the 594-bp upstream intergenic region) using primer sets listed in Table 2. Three overlapping DNA fragments, designated P*bcct1* probe A (120-bp), probe B (110-bp), and probe C (101-bp), were generated from the *bcct1* regulatory region (includes 15-bp of the coding region and the 276-bp upstream intergenic region) using primer sets listed in Table 2. Two overlapping DNA fragments, designated P*bcct3* probe A (108-bp) and probe B (107-bp), were generated from the *bcct3* regulatory region (includes 17 bp of the coding region and 179-bp of the upstream intergenic region) using primer sets listed in Table 2. Four overlapping DNA fragments, designated P*proV1* probe A (160-bp), probe B (134-bp), probe C (108-bp), and probe D (109-bp), were generated from the *proV1* regulatory region (includes 9 bp of the coding region and the 438-bp upstream intergenic region) using primer sets listed in Table 2. Fragments designated P*bcct2* (233-bp) and P*bcct4* (244-bp) were generated from the *bcct2* and *bcct4* regulatory regions, respectively, using primers listed in Table 2. Two overlapping DNA fragments, designated P*cosR* probe A (105-bp) and probe B (142-bp), were generated from the *cosR* regulatory region (includes 4-bp of the coding region and 216 bp of the upstream intergenic region) using primer sets listed in Table 2. The concentration of purified CosR-His and OpaR was determined using a Bradford assay. CosR or OpaR was incubated for 20 minutes with 30 ng of each DNA fragment in a defined binding buffer (10 mM Tris, 150 mM KCl, 0.5 mM dithiothreitol, 0.1 mM EDTA, 5% polyethylene glycol [PEG] [pH 7.9 at 4°C]). A 6% native acrylamide gel was pre-run for 2 hours at 4C (200 V) in 1 X TAE buffer. Gels were loaded with the DNA:protein mixtures (10 μL), and run for 2 hours at 4°C (200 V). Finally, gels were stained in an ethidium bromide bath for 15 min and imaged.

### Reporter Assays

A GFP reporter assay was conducted using the *E. coli* strain MKH13 (69). GFP reporter plasmids were constructed as previously described (17). Briefly, each regulatory region of interest was amplified using primers listed in Table 2 and ligated via Gibson assembly protocol with the promoterless parent vector pRU1064, which had been digested with SpeI, to generate reporter plasmids with GFP under the control of the regulatory region of interest. Complementary regions for Gibson assembly are indicated in lower case letters in the primer sequence (Table 2). Reporter plasmid P*_betI_*-*gfp* encompasses 594-bp upstream of the *betIBAproXWV* operon. Reporter plasmid P*_bcct1_*-*gfp* encompasses 278-bp upstream of the P*bcct1* regulatory region. Reporter plasmid P*_bcct3_*-*gfp* encompasses 397-bp upstream of the P*bcct3* regulatory region. Reporter plasmid P*_proV1_*-*gfp* encompasses 438-bp upstream of the P*proV1* regulatory region. Reporter plasmid P*_cosR_*-*gfp* encompasses 397-bp upstream of the P*cosR* regulatory region. The full-length *cosR* was then expressed from an IPTG-inducible promoter in the pBBR1MCS expression vector. Relative fluorescence (RFU) and OD_595_ were measured; specific fluorescence was calculated by dividing RFU by OD_595_. Strains were grown overnight with aeration at 37°C in LB1% with ampicillin (50 µg/mL) and chloramphenicol (12.5 µg/mL), washed twice with 1X PBS, then diluted 1:1000 in M9G1%. Expression of *cosR* was induced with 0.25 mM IPTG, and strains were grown for 20 hours at 37°C with aeration under antibiotic selection. GFP fluorescence was measured with excitation at 385 and emission at 509 nm in black, clear-bottom 96-well plates on a Tecan Spark microplate reader with Magellan software (Tecan Systems Inc., San Jose, CA). Specific fluorescence was calculated for each sampled by normalizing fluorescence intensity to OD_595_. Two biological replicates were performed for each assay.

A GFP reporter assay was conducted in RIMD2210633 wild-type, Δ*betI* and Δ*opaR* mutant strains. The P*_betI_*-*gfp* reporter plasmid was transformed into *E. coli* β2155 λ*pir* and conjugated into wild-type, Δ*betI* and Δ*opaR* mutant strains. Strains were grown overnight with aeration at 37°C in LB3% with tetracycline (1 µg/mL). Cells were then pelleted, washed two times with 1X PBS, diluted 1:100 into M9G3% and grown for 20 hours with antibiotic selection. Choline was added to a final concentration of 1 mM, where indicated. GFP fluorescence was measured with excitation at 385 and emission at 509 nm in black, clear-bottom 96-well plates on a Tecan Spark microplate reader with Magellan software (Tecan Systems Inc.). Specific fluorescence was calculated for each sampled by normalizing fluorescence intensity to OD_595_. Two biological replicates were performed for each assay.

### Bioinformatics analysis

The *V. parahaemolyticus* protein CosR (NP_798285) was used as a seed for BLASTp to identify homologs in the *Vibrionaceae* family in the NCBI database. Sequences of representative strains were downloaded from NCBI and used in a Python-based program Easyfig to visualize gene arrangements (70). Accession numbers for select strains were: *V. parahaemolyticus* RIMD (BA00031), *V. crassotreae* 9CS106 (CP016229), *V. splendidus* BST398 (CP031056), *V. celticus* CECT7224 (NZ_FLQZ01000088), *V. lentus* 10N.286.51.B9 (NZ_MCUE01000044), *V. tasmaniensis* LGP32 (FM954973), *V. cyclitrophicus* ECSMB14105 (CO039701), *Aliivibrio fischeri* ES114 (CP000021), *A. fischeri* MJ11 (CP001133), *A. wodanis* AWOD1 (LN554847), *A. wodanis* 06/09/160 (CP039701). The *V. parahaemolyticus* RIMD2201633 CosR and *A. fischeri* ES114 EctR protein sequences were retrieved from NCBI using accession numbers NP_798285 and AAW88191.1, respectively, and input into the SWISS-MODEL workspace, which generated a 3D model of a homodimer to identify putative ligand-binding pockets (71–75).

## Acknowledgements

This research was supported by a National Science Foundation grant (award IOS-1656688) to E.F.B. G.J.G. was funded in part by a University of Delaware graduate fellowship award. DPM was supported by a departmental undergraduate researcher fellowship. We thank members of the Boyd Group for constructive feedback on the manuscript.

